# Short- and long-term effects of uterine disease on oocyte developmental capacity in postpartum dairy cows

**DOI:** 10.1101/2025.02.07.636469

**Authors:** M. O. Caldeira, K. S. McDonald, E. S. M. Martinez, J. G. N. Moraes, I. Sellmer Ramos, S. E. Poock, M. S. Ortega, M. C. Lucy

## Abstract

The hypothesis was that early postpartum uterine disease would reduce the developmental capacity of oocytes thus contributing to the reduced fertility of dairy cows with uterine disease. Dairy cows were diagnosed healthy or with metritis at 7 to 10 d postpartum. The reproductive tract was collected at approximately 1 mo (Exp. 1) or approximately 80 or 165 d (Exp. 2) postpartum for the collection of cumulus-oocyte complexes (COC). The COC were matured, co-incubated with sperm for fertilization, and cultured to the blastocyst stage (8 d) in vitro. For Exp.1, the disease diagnosis (healthy or metritis) did not affect the number of collected COC or the subsequent embryo development to the blastocyst stage. The presence of purulent material in the uterine lumen (endometritis) at time of oocyte collection, however, was associated with a reduced cleavage rate evaluated 3 d following fertilization. For Exp. 2, there was no effect of disease diagnosis (healthy or metritis) on the number of COC or their subsequent development. Reduced cleavage rates were observed in COC retrieved from cows slaughtered at 80 d postpartum, but not at 165 d postpartum, and this reduction was associated with a vaginal microbiome indicative of uterine disease at 4 to 5 wk postpartum. Regression analyses that included plasma haptoglobin or energy metabolite concentrations or uterine bacterial genera abundance did not explain a large percentage of the variation in oocyte development in vitro. We conclude that there is an effect of uterine disease at one month postpartum on the oocyte and its capacity for development (Exp. 1) and this effect may be present at 80 d postpartum (Exp. 2). In later postpartum cows (165 d postpartum; Exp. 2) there was no effect of uterine disease on in vitro oocyte development.

**GRAPHICAL ABSTRACT:** 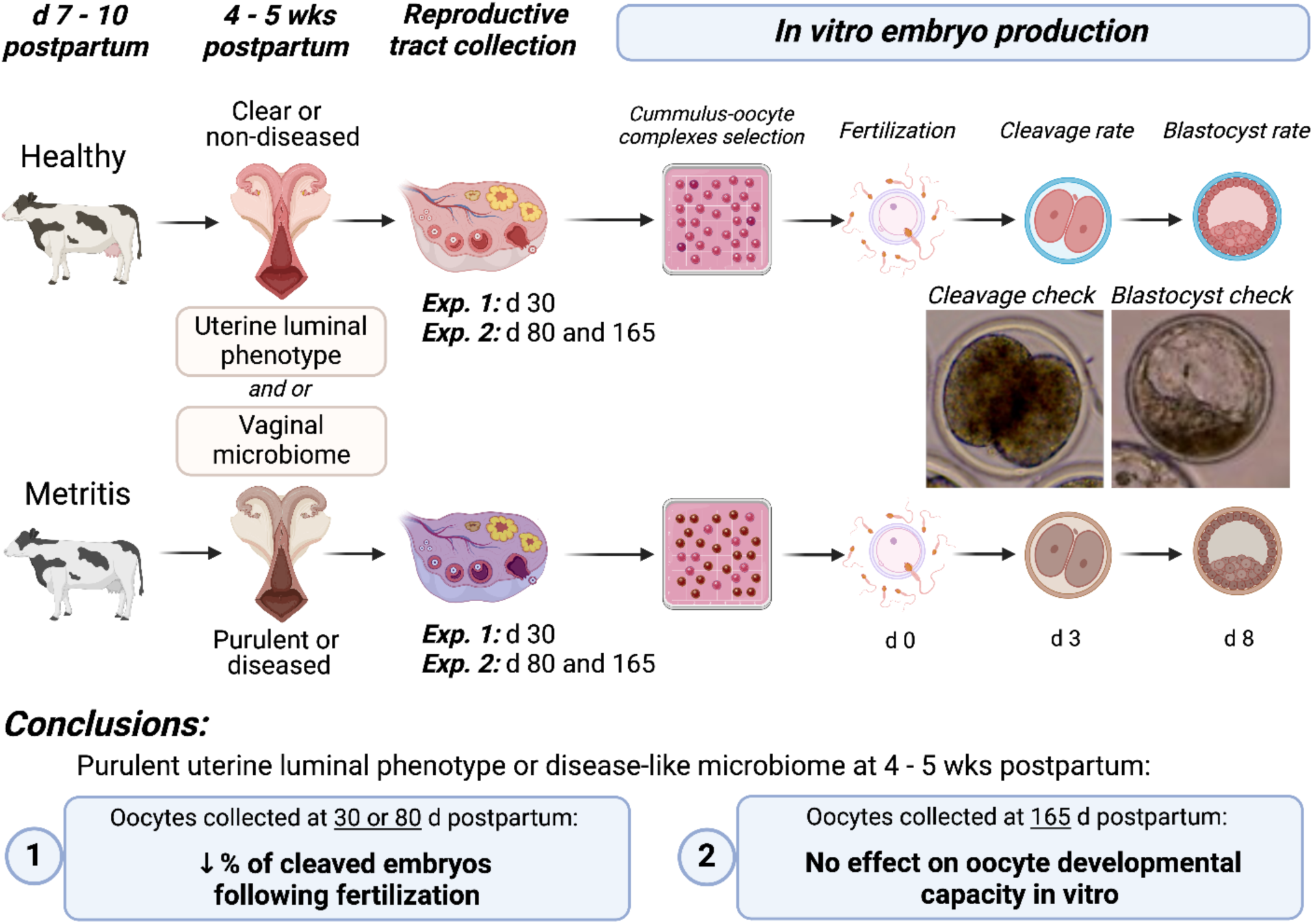

**HIGHLIGHTS:**

- Oocytes collected at 1 month postpartum from dairy cows with endometritis (purulent uterine lumen) had a lower cleavage rate following in vitro fertilization when compared with oocytes collected from healthy cows (Exp. 1).
- Oocytes collected at approximately 80 d postpartum from dairy cows with evidence of uterine disease at 4 to 5 wk postpartum had a lower cleavage rate following in vitro fertilization but this disease-associated difference was not observed when oocytes were collected from cows later postpartum (approximately 165 d postpartum; Exp. 2).
- Regardless of the study (Exp. 1 or 2), uterine disease primarily affected the percentage of oocytes that cleaved after fertilization.
- Statistical associations between circulating metabolites or relative abundance of uterine bacteria were either not significant or explained only a small percentage of the variation in the in vitro embryo development in either experiment.

## 1. Introduction

Postpartum metritis in dairy cows is characterized by an enlarged uterus and fetid red-brown or purulent uterine discharge that is typically diagnosed within 7 to 10 d postpartum. The expectation is that 10 to 20% of dairy cows will develop metritis after calving [1–3]. Most cows recover from metritis within the first month postpartum. Cows with metritis are at an increased risk of developing chronic uterine inflammation (endometritis), which can have long-term negative effects on fertility [4,5]. In addition to uterine disease, postpartum cows may experience negative energy balance and associated metabolic disease postpartum that can affect fertility [6,7]. One current hypothesis for infertility in postpartum cows is that the combined effects of uterine and metabolic disease have a negative effect on oocyte quality leading to defects in fertilization or early embryonic development [2,6–8].

Most studies testing the effects of inflammatory molecules or metabolites use oocytes removed from healthy cows and treated in vitro. Fewer studies have performed similar work on the whole animal. Dickson et al. [9] experimentally induced endometritis in nonlactating dairy cows and reported a reduction in embryo development to the morula stage following in vitro fertilization of aspirated oocytes. Ribeiro et al. [10] flushed embryos from postpartum cows and reported a reduction in cleaved, live, and high-quality embryos in cows that previously experienced uterine disease.

In this work, the results of the previous studies [9,10] were extended by specifically working with primiparous postpartum cows with or without metritis, and COC were collected for in vitro embryo production (IVP) during three timepoints: 30, 80, 165 d after parturition. Collectively, this research examines the short-term (oocytes collected at approximately 1 month postpartum) and long-term (oocytes collected at approximately 80 to 165 d postpartum) effects of early postpartum uterine disease on oocyte developmental capacity. The current study is different from all other previously published work because we worked with early postpartum lactating cows and removed oocytes from cows to separate the biological effect of the oviductal and uterine environment from the capacity of the oocyte to develop. We hypothesized that uterine disease would negatively affect oocyte developmental competence. We also measured the relative importance of systemic factors by measuring an acute phase protein (haptoglobin) and energy metabolites [non-esterified fatty acids (NEFA), beta-hydroxybutyrate (BHB), and glucose] and associating their plasma concentrations postpartum with subsequent developmental competence of the oocyte. Finally, we tested the association between oocyte development and bacterial genera present in the reproductive tract.

## 2. Materials and methods

### 2.1 Ethics statement

Study procedures were approved by the University of Missouri (MU) Institutional Animal Care and Use Committee (Protocol number 9635).

### 2.2 Study design

For Exp. 1 and 2, first parity Holstein cows were selected at clinical disease diagnosis (7 to 10 d postpartum) from a confinement herd in eastern Kansas or the MU herd. Cows with a clinical diagnosis of metritis (fetid red-brown watery vaginal discharge with a flaccid uterus) were selected and matched with clinically healthy cows [viscous (not watery) and non-fetid discharge] that calved during the same wk. Detailed methods for Exp. 1 and 2 are published [11,12]. A brief description is presented in the next two paragraphs.

For Exp. 1, there were 18 metritis cows and 17 healthy control cows. Cows were either antibiotic-treated [metritis-treated (n=9) and healthy-treated (n=9)] with ceftiofur hydrochloride (i.m. 1.25 g/d for 3 d; Excenel RTU, Zoetis, Parsippany-Troy Hills, NJ, USA) or untreated [metritis-untreated (n=9) and healthy-untreated (n=8)]. For cows coming from Kansas, diagnosis was performed within the Kansas herd and cows were moved on the same day to the MU herd where antibiotic treatment was administered, and they remained until the end of the experiment. Cows were humanely slaughtered by captive bolt stunning and exsanguination at 29.1 ± 1.7 d postpartum (approximately 1 mo postpartum).

For Exp. 2, all cows were selected from the Kansas herd used in Exp. 1, but stayed in the Kansas herd until 1 d prior to tissue collection following slaughter at approximately 80 d postpartum (5 metritis and 5 control; 79.0 ± 7.5 d postpartum; mean ± SD; 80 d postpartum group) or approximately 165 d postpartum (5 metritis and 4 control; 165.0 ± 4.9 d postpartum; mean ± SD; 165 d postpartum group). Preliminary results from Exp. 1 demonstrated no long-term effect of antibiotic treatment on a variety of study endpoints including microbiome [11]. Cows in Exp. 2, therefore, were treated with ceftiofur hydrochloride at the discretion of the herdsman. Cows in Exp. 2 were treated with a sequence of two PGF2α injections at a 14 d interval prior to slaughter (d -32 PGF2α; d -18 PGF2α) to target the luteal phase at slaughter.

### 2.3 Analyses of uterine and vaginal microbiome

Detailed procedures for the collection and analysis of uterine and vaginal microbiome from Exp. 1 and 2 are published [11,12]. A brief description is presented in this section. For Exp. 1, cows had their uterine contents sampled for 16S rRNA gene sequencing when they were identified and selected for the trial. Uterine swabs were collected transcervically using a double-guarded culture swab (Jorgensen Laboratories, Loveland, CO) following perineal cleaning and disinfection. Swab samples were carefully moved to a sterile cryovial (CryoTube Vial; Thermo Fisher Scientific; Waltham, MA), immediately placed in dry ice, and subsequently stored in a - 80°C freezer. Promptly after uterine swab collection, cows were either treated with antibiotics or were left untreated as described in Section 2.2. Following slaughter (1 mo postpartum), the uterus was removed from the abdomen and taken to the Veterinary Medical Diagnostic Laboratory at the College of Veterinary Medicine at MU. The uterus and ovaries were placed inside a biosafety cabinet. The outside of the uterus was then cleaned and disinfected and the uterine lumen exposed by dissection. A sample of the uterine lumen of the previously gravid and non-gravid uterine horns was collected and placed in a CryoTube and frozen in liquid nitrogen to be used for 16S rRNA DNA sequencing. The samples were stored in a -80°C freezer prior to sequencing.

For Exp. 2, the vaginal microbiome was sampled weekly from enrollment until 35 ± 2 d postpartum, using a double-guarded uterine swab (Jorgensen Laboratories). The perineal area was cleaned with a paper towel, disinfected with 70% ethanol and dried with a clean paper towel. The swab was introduced cranially in the vagina to reach the cervical entrance and rolled against the mucosa to obtain the sample. Swabs were frozen on dry ice and transported to the laboratory where they were stored at -80°C prior to metagenomic sequencing. The rationale behind the use of vaginal swabs was that the cervix was relatively open and uterine contents would exude through the cervix and into the vagina. Previous studies have demonstrated an association between the vaginal and uterine microbiota in early postpartum cows with uterine disease [13].

Swab samples (Exp. 1 uterine sample at disease diagnosis; Exp. 2, vaginal samples at wk 1 to 5) and uterine luminal tissue samples (Exp. 1 at one month postpartum) were subjected to 16S rRNA gene sequencing. The DNA extraction, library construction and sequencing were performed at the Metagenomics Center and the Genomics Technology Core of the University of Missouri. Amplicon sequences were processed and analyzed using QIIME2 (version 2020.6, https://qiime2.org) as previously described [11,12]. Amplicon sequence variants (ASV) sharing the same taxa were collapsed together at the species level using QIIME2 taxa collapse function and further collapsed to the genus level using PROC SUMMARY of SAS 9.4 (SAS Institute Inc., Cary, NC). For Exp. 2, The ASV read counts data from vaginal swab samples collected during the first and second wk postpartum were added together within individual cows and defined as the ASV at “disease diagnosis”. Likewise, the ASV read counts from the fourth and fifth wk were added together and defined as “4 to 5 wk postpartum”. The rationale for combining the sample d into a single time point was to create an integrated sample that encompassed a longer time that bracketed data around the time of disease diagnosis as well as several wk later.

The relative abundance of a microorganism was calculated within cow using the equation: relative abundance = number of ASV counts of each microorganism/sum of all counts in that individual cow, expressed as a percentage. The microbiome of the uterus (Exp. 1) and vagina (Exp. 2) differed for healthy versus diseased cows at disease diagnosis (7 to 10 d postpartum) [11,12].

Exp. 1 included the measurement of the microbiome from tissues collected from cows at 1 mo postpartum. This was not possible for cows in Exp. 2 whose reproductive tracts would be collected later postpartum. To group cows based on their uterine disease status in Exp. 2, principal components (**PC**) were fitted for each cow based on their respective vaginal microbiomes at 4 to 5 wk postpartum). The PC were used to identify groups of cows with microbiome that were consistent with a diseased or non-diseased uterus (procedures described in detail [12]). This disease classification (based on PC arising from the microbiome) was independent of the original diagnosis on farm.

### 2.4 Blood collection and analysis

Blood samples used in these analyses were collected thrice weekly for Exp. 1 (first 30 d) and weekly (first 5 wk) for Exp. 2. Samples were collected from the coccygeal veins (10 ml Monoject EDTA, Covidien, Mansfield MA) after morning milking and typically when cows were being fed. Plasma was separated by centrifugation (2,000 ×*g* for 15 min at 4°C) and stored at -20°C. Plasma haptoglobin concentrations were measured with a commercial ELISA kit (Immunology Consultants Laboratory, Inc.; catalog number E-10HTP, Portland, OR) following the manufacturer’s protocol. Plasma NEFA concentrations were determined using the NEFA HR (2) commercial kit (Wako Diagnostics, Richmond, VA). Glucose concentrations were measured enzymatically using the glucose oxidase method (Pointe Scientific Inc., Canton, MI). β- hydroxybutyrate concentrations were determined by enzymatic assay (RB1007; Randox Laboratories Ltd., Antrim, UK). Absorbance was quantified using a microplate reader spectrophotometer (Synergy 2 BioTek, Winooski, VT). Plasma progesterone concentrations prior to slaughter were measured using a validated radioimmunoassay[14]. In Exp. 1, there were 3 out of 17 metritic cows and 7 out of 18 healthy cows cycling (> 1 ng/mL progesterone) prior to slaughter. All cows in Exp. 2 were cycling before slaughter with an average interval to cyclicity of 36.2 ± 14.3 d postpartum (mean ± SD).

### 2.4 Evaluation of the uterine contents

The reproductive tract was removed from the abdomen following slaughter, processed for the collection of microbiome samples at the MU Veterinary Medical Diagnostic Laboratory and then transported to a second laboratory in the Animal Science building at MU where the uterine lumen was flushed with sterile saline. The uterine luminal fluid was classified as either containing purulent material (purulent flush; containing white, yellow, or sanguineous purulent material) or clear (clear flush; translucid and clear mucus without pus). In Exp.1 there were 15 cows with a purulent flush and 20 cows with a clear flush. All the Exp. 2 cows had a clear uterine flush. The total time from slaughter to processing in the Animal Science laboratory was 1 to 2 h.

### 2.5 In vitro maturation, fertilization, and culture

The IVF and in vitro culture were conducted similarly for Exp. 1 and 2. The ovaries were trimmed from the tracts and stayed in saline solution at room temperature until oocyte collection. All media [oocyte collection (**OCM**), oocyte maturation (**OMM**), fertilization (**IVF-TALP**), wash (**HEPES-TALP**), culture (**SOF-BE2**), and the supplement penicillamine, hypotaurine, and epinephrine (**PHE**)] were produced in house following previously described protocols [15]. The only ready-to-use material was the sperm purification gradient (Isolate®, Irvine Scientific, Santa Ana, CA, USA). The surface of each ovary was cut with a scalpel to harvest immature cumulus-oocyte complexes (COC) from follicles 2 to 8 mm in diameter into OCM. After harvesting, COC were washed in OCM and those surrounded by at least two layers of cumulus cells and with an evenly granulated cytoplasm were used. COC recovered from each cow were maintained as an individual replica throughout culture, maturation, and fertilization. After 24 h maturation, fertilization was performed in vitro according to the methods described by [15] and [16]. Briefly, matured COC from each cow were washed three times in HEPES-TALP and placed in a 35 mm dish containing 1.7 ml of IVF-TALP. Sperm from a high-fertility Holstein bull was purified from a frozen-thawed straw using an isolate gradient, diluted with IVF-TALP to a final concentration of 1 × 10^6^/mL in the dish, and supplemented with PHE to increase sperm motility and capacitation. Fertilization occurred by co-incubation of sperm and COC for 18 to 22 h in a humidified atmosphere at 38.5°C and 5% (v/v) CO2. At the end of fertilization, putative zygotes (oocytes exposed to sperm) were denuded from surrounding cumulus cells by vortexing for 5 min in 600 μL of HEPES-TALP containing 10,000 U/mL of hyaluronidase. Embryos were then cultured in four-well dishes in SOF-BE2 covered with mineral oil at 38.5°C in a humidified atmosphere of 5% (v/v) O2 and 5% (v/v) CO2 with the balance N2 until collection. Day 0 was considered the day of insemination. The percentage of putative zygotes cleaved was recorded on d 3, and the blastocyst rate on d 8 post-insemination.

### 2.6 Statistical analysis

The primary purpose was to evaluate the effect of uterine disease on oocyte developmental competence. The dependent variables were: 1) number of COC collected per cow; 2) percentage of cleaved embryos following fertilization [100*(number of cleaved embryos/number of putative zygotes placed in culture); 3) percentage blastocysts formed expressed as a function of the number of putative zygotes placed in culture; and 4) percentage of blastocysts formed expressed as a function of cleaved embryos. All data were analyzed using the mixed model procedure (PROC MIXED) of SAS 9.4. For Exp. 1, a full model that included status at disease diagnosis (healthy or metritis), treatment (antibiotic or control), uterine flush (clear or purulent), cyclicity and all interactions was initially fit. The effect of antibiotic treatment was not significant (*P >* 0.10) so a reduced model (status, flush, cyclicity and interactions) was tested. Cow nested within status by flush by cyclicity was included as a random effect. For Exp. 2, all cows were cycling prior to slaughter and all flushes were clear so the effects of cyclicity and flush phenotype were not tested. The model for Exp. 2 included status at disease diagnosis (healthy or metritis), day of oocyte collection (d 80 or 165) and their interactions. Cow nested within status and day was included as a random effect. This model yielded no significant effects, so a second model was tested that included the uterine disease status at 4 to 5 wk postpartum. Methodology used to assign a uterine disease status based on vaginal microbiome at 4 to 5 wk postpartum was previously described [12]. This second model included disease status (non-diseased or diseased), day of oocyte collection (d 80 or 165) and interactions. Cow nested within status and day was included as a random effect. In a final set of analyses for Exp. 1 and 2, circulating concentrations of haptoglobin, glucose, NEFA, and BHB and the abundance of bacterial genera within the uterine (Exp. 1) or vaginal (Exp. 2) microbiome postpartum were tested for their effects on oocyte development using linear regression (PROC REG of SAS with the stepwise model selection procedure). The PROC REG with stepwise selection is designed to build a predictive linear model without overfitting (including too many independent variables). In stepwise selection variables enter the model but may not necessarily remain in the final model. The dependent variables were number of COC, percentage of cleaved embryos, etc. The regressors were the average concentrations of haptoglobin, glucose, NEFA, and BHB during the first 15 d postpartum (early) or between 16 and 30 d postpartum (late) and the relative abundance of bacterial genera within the uterus at one month postpartum (Exp. 1) or the vaginal microbiome at 4 to 5 wk postpartum (Exp. 2). The cutoff for entry into the model was *P <* 0.15 and the cutoff to stay in the model was P < 0.05. Across all analyses, significance was declared at *P <* 0.05 and a statistical tendency was 0.05 *< P <* 0.10. Data are presented as lsmeans ± SEM unless stated otherwise.

## 3 Results

### 3.1 In vitro embryo development (Exp. 1; oocytes collected at 1 month postpartum)

There was no effect of the original disease diagnosis (healthy or metritis; diagnosed at 7 to 10 d postpartum) on the number of COC collected (*P =* 0.806; **Figure 1A**), the percentage of cleaved embryos (*P =* 0.776; **Figure 1B**), or the percentage of cleaved embryos or embryos developing to blastocysts (*P =* 0.613 and *P =* 0.233, respectively; **Figures 1C and 1D**). The number of COC collected (**Figure 1A**) and the percentage of cleaved embryos or embryos developing to blastocysts (**Figures 1C and 1D**) were also similar for cows that had a clear versus purulent flush at the time of oocyte collection (1 mo postpartum). The uterine flush phenotype at one month, however, was associated with the percentage of cleaved embryos (**Figure 1B**; *P =* 0.017). Cows with a clear uterine flush had a greater percentage of cleaved embryos (69.9 ± 6.2%) compared with cows with a purulent uterine flush (46.3 ± 7.0%).

**Figure 1.**
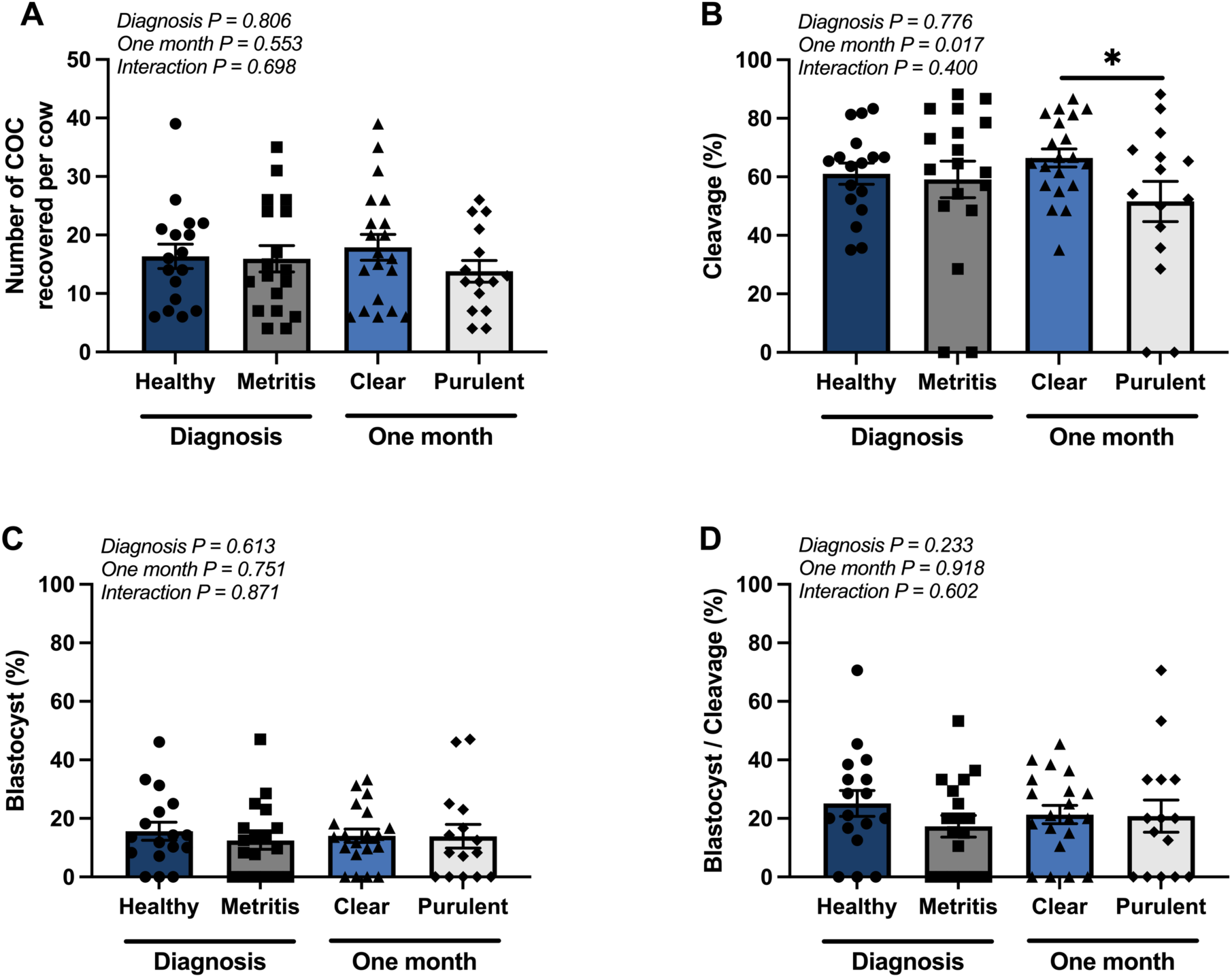
Effects of uterine disease on oocyte developmental capacity following oocyte collection and *in vitro* fertilization at one month postpartum (Exp. 1). Cows were diagnosed as Healthy (normal uterine discharge; n=17) or Metritis (fetid red-brown watery uterine discharge; n=18) within 7 to 10 d postpartum. Cows had their uterine lumen flushed during tissue collection at 1 mo postpartum and were classified based on their flush phenotype as either clear (n=20) or containing purulent material (n=15). Each dot represents an individual cow. The number of cumulus-oocyte complexes (COC) recovered per cow (A), the percentage of embryos cleaved by d 3 (cleavage percentage; B), percentage blastocysts on d 8 post insemination as a function of COC (blastocyst percentage; C), and the percentage of blastocysts as a function of cleaved embryos (blastocyst/cleavage percentage; D) are shown. P-values from the statistical analysis for the effect of disease at 7 to 10 d postpartum (diagnosis) and at 1 mo postpartum (clear or purulent flush) are provided for each individual graph. Significance was declared when *P <* 0.05 and is represented by an asterisk.

There was an effect of cyclicity status on the number of COC and a tendency for an interaction of cyclicity status with flush phenotype at one month postpartum for the percentage of cleaved embryos. There were fewer (*P <* 0.039) COC collected from cows that were cycling (9.8 ± 3.5 oocytes/cow) compared with those that were non-cycling (18.3 ± 1.8 oocytes per cow) by the time of oocyte collection (1 mo postpartum). The greatest percentage of cleaved embryos was observed in the cycling cows with a clear flush and the least percentage of cleaved embryos was observed in the cycling cows with a purulent flush (flush phenotype by cyclicity interaction (*P <* 0.066; 61.4 ± 5.7, 78.5 ± 11.0, 55.6 ± 6.2, and 37.1 ± 12.4 for clear-noncycling, clear- cycling, purulent-noncycling and purulent cycling, respectively).

### 3.2 In vitro embryo development (Exp. 2; oocytes collected at approximately 80 and 165 d postpartum)

There was no effect of the original disease diagnosis (healthy or metritis; diagnosed at 7 to 10 d postpartum) on the number of COC collected per cow (9.7 ± 1.9 versus 9.3 ± 1.8; *P =* 0.894), the percentage of cleaved embryos (56.0 ± 11.5 versus 44.2 ± 10.8; *P =* 0.464), the percentage of embryos developing to blastocyst (17.8 ± 6.7 versus 16.4 ± 6.3; *P =* 0.879), or cleaved embryos developing to blastocysts (27.6 ± 9.6 versus 23.0 ± 9.4; *P =* 0.746) (healthy versus metritis, respectively). Interactions of disease diagnosis with day were not detected (*P >* 0.10) for any of the dependent variables (COC, percentage of cleaved embryos, etc.). There was a tendency (*P =* 0.097) for an effect of day on the number of COC collected per cow with a greater number of COC collected from the later-postpartum cows (7.2 ± 1.8 versus 11.8 ± 1.9).

The model that included the original disease diagnosis yielded no significant effects (previous paragraph) so a second model that included the uterine disease status at 4 to 5 wk postpartum was tested. The methodology used to assign a uterine disease status based on vaginal microbiome at 4 to 5 wk postpartum were described [12]. When the uterine status at 4 to 5 wk (non-diseased versus diseased) was used, there was no effect of disease status on the number of COC collected per cow (9.5 ± 2.0 versus 9.6 ± 1.8; *P =* 0.976; **Figure 2A**), the percentage of embryos developing to blastocyst (16.3 ± 7.0 versus 15.8 ± 6.4; *P =* 0.959; **Figure 2C**), or cleaved embryos developing to blastocysts (23.1 ± 10.7 versus 25.4 ± 9.8; *P =* 0.874; **Figure 2D**) (non-diseased versus diseased, respectively). Neither an effect of day nor an interaction of disease with day were detected (*P >* 0.10) for the number of COC (F**igure 2A**), the percentage of embryos developing to blastocyst (**Figure 2C**), or cleaved embryos developing to blastocysts (**Figure 2D**). There was a disease by day interaction for the percentage of cleaved embryos (**Figure 2B**). The percentage of cleaved embryos was less (*P <* 0.0195) for diseased versus non-diseased cows when oocytes were collected on approximately d 80 (25.9 ± 14.5 versus 74.7 ± 11.8; diseased versus non-diseased; **Figure 2C**). Later postpartum (d 165), the percentage of cleaved embryos was similar (*P =* 0.4126) for diseased (50.7 ± 11.8) and non-diseased (33.5 ± 16.7).

**Figure 2.**
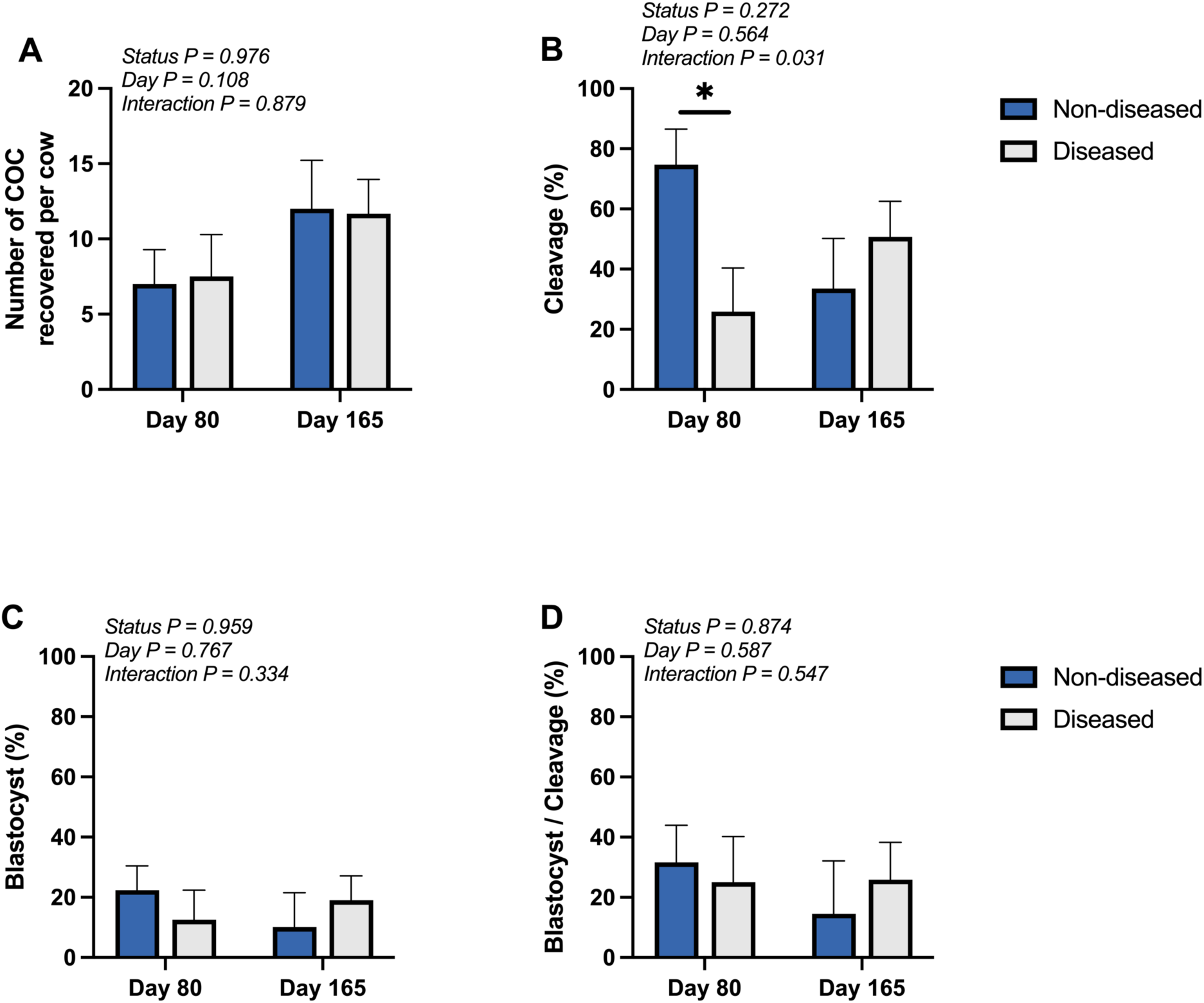
Effects of uterine disease on oocyte developmental capacity following oocyte collection and *in vitro* fertilization at 80 to 165 d postpartum (Exp. 2). Cows were classified as non-diseased or diseased based on the analysis of principal components of the vaginal microbiome at 4 to 5 wk postpartum and slaughtered at approximately 80 or 165 d postpartum creating 4 groups of cows [non-diseased d 80 (n=6), diseased d 80 (n=4), non-diseased d 165 (n=3) and diseased d 165 (n=6)] [12]. Each dot represents an individual cow. The number of cumulus-oocyte complexes (COC) recovered per cow (A), the percentage of cleaved embryos by d 3 (cleavage percentage; B), percentage blastocysts on d 8 post insemination as a function of COC (blastocyst percentage; C), and the percentage of blastocysts as a function of cleaved embryos (blastocyst/cleavage percentage; D) are shown. P-values from the statistical analysis for the effect of disease status at 4 to 5 wk postpartum and day are provided for each individual graph. Significance was declared when *P <* 0.05 and is represented by an asterisk.

### 3.3 Predictive modeling of individual cow factors affecting in vitro oocyte development for oocytes collected at 1 month postpartum (Exp. 1)

Plasma concentrations of haptoglobin, glucose, NEFA, and BHB during the first 15 d postpartum (early) and from 16 to 30 d postpartum (late) are presented for cows based on their initial disease diagnosis and based on the uterine flush phenotype at the time of slaughter (**Table 1**). As expected, there was an effect (*P <* 0.001) of the initial disease diagnosis on plasma haptoglobin concentrations during the early period (greater in metritis cows; **Table 1**). Glucose (*P =* 0.664), NEFA (*P =* 0.905), and BHB (*P =* 0.110) were similar for healthy versus metritis during the early period. During the late period, plasma haptoglobin concentrations had decreased in metritis cows and were not different from healthy (**Table 1**). Plasma glucose concentrations were similar, but plasma NEFA and BHB concentrations were greater in healthy compared with metritis cows (**Table 1**).

**Table 1.** Plasma haptoglobin, glucose, non-esterified fatty acid (NEFA), and β-hydroxybutyrate (BHB) concentrations for Exp. 1. Cows were diagnosed as healthy or metritis at 7 to 10 d postpartum. The uterine lumen was flushed with PBS and classified as clear or containing purulent material at slaughter. Data are presented as mean ± SEM.

|  | Initial disease diagnosis |  |  | Uterine flush (1 mo postpartum) |  |  |
| --- | --- | --- | --- | --- | --- | --- |
|  | Healthy<br>(n=17) | Metritis<br>(n=18) | *P*-value | Clear (n=20) | Purulent<br>(n=15) | *P*-value |
| Early<sup>1</sup> |  |  |  |  |  |  |
| Haptoglobin (g/L) | 0.2 ± 0.2 | 1.5 ± 0.2 | <0.001 | 0.7 ± 0.2 | 1.2 ± 0.3 | ns |
| Glucose (mg/dL) | 67.8 ± 1.9 | 66.7 ± 1.8 | ns<sup>3</sup> | 64.9 ± 1.6 | 70.3 ± 1.9 | 0.036 |
| NEFA (μEq/L) | 428.9 ± 43.2 | 436.1 ± 42.0 | ns | 413.7 ± 39.5 | 457.8 ± 45.6 | ns |
| BHB (mmol/L) | 0.67 ± 0.04 | 0.76 ± 0.04 | ns | 0.72 ± 0.04 | 0.71 ± 0.05 | ns |
| Late<sup>2</sup> |  |  |  |  |  |  |
| Haptoglobin (g/L) | 0.2 ± 0.1 | 0.3 ± 0.1 | ns | 0.2 ± 0.1 | 0.3 ± 0.1 | ns |
| Glucose (mg/dL) | 66.4 ± 1.8 | 64.8 ± 1.7 | ns | 66.1 ± 1.6 | 64.9 ± 1.9 | ns |
| NEFA (μEq/L) | 424.0 ± 33.8 | 299.3 ± 32.9 | 0.012 | 391.5 ± 33.3 | 317.7 ± 38.4 | ns |
| BHB (mmol/L) | 0.81 ± 0.04 | 0.68 ± 0.04 | 0.019 | 0.79 ± 0.04 | 0.67 ± 0.04 | 0.031 |

<sup>1</sup>Average of all samples from disease diagnosis to 15 d postpartum.
<sup>2</sup>Average of all samples from 16 d postpartum until tissue collection at 1 mo postpartum.
<sup>3</sup>ns = not significant

Plasma concentrations of haptoglobin (*P =* 0.153 and *P =* 0.854) and NEFA (*P =* 0.471 and *P =* 0.156) were similar for clear versus purulent cows regardless of the sample period (early and late; respectively; **Table 1**). Plasma glucose concentrations were less (early period; *P <* 0.036) and plasma BHB concentrations were greater (late period; *P <* 0.031) in cows with a clear compared with purulent flush (**Table 1**).

The average plasma concentrations of haptoglobin, glucose, NEFA, and BHB during the early and late periods as well as the relative abundance of the 10 most-abundant genera sequenced from the uterine lumen of individual cows at one month postpartum were included in a stepwise regression analysis (PROC REG of SAS) to build a predictive model for the number of COC collected per cow, the percentage of cleaved embryos, the percentage of embryos developing to blastocysts, and the percentage of cleaved embryos developing to blastocysts (**Table 2**). Beginning with the 18 individual regressors, the stepwise regression retained a single regressor for the number of COC (*Cutibacterium* abundance; positively associated with the number of COC), a single regressor for cleavage percentage (*Fusobacterium* abundance; negatively associated with cleavage percentage) and two regressors for the percentage of blastocysts developing from cleaved embryos [*Mycoplasma* abundance (negatively associated) and *Cutibacterium* abundance (positively associated)] (**Table 3**). None of the regressors were significant for the percentage of blastocysts developing from putative zygotes. The regression procedure did not retain any of the regressors for early or late metabolite concentrations.

**Table 2.** Statistical parameters for individual regressor variables included in the PROC REG model for the prediction of oocyte developmental competence (Exp. 1).

|  | N | Mean | SD | Min | Max |
| --- | --- | --- | --- | --- | --- |
| Early<sup>1</sup> |  |  |  |  |  |
| Haptoglobin (g/L) | 35 | 0.88 | 1.00 | 0.00 | 3.43 |
| Glucose (mg/dL) | 35 | 67.19 | 7.61 | 50.28 | 81.65 |
| NEFA (μEq/L) | 35 | 432.60 | 175.58 | 121.40 | 926.25 |
| BHB (mmol/L) | 35 | 0.72 | 0.18 | 0.43 | 1.20 |
| Late<sup>2</sup> |  |  |  |  |  |
| Haptoglobin (g/L) | 35 | 0.25 | 0.51 | 0.00 | 2.23 |
| Glucose (mg/dL) | 35 | 65.60 | 7.22 | 52.62 | 86.45 |
| NEFA (μEq/L) | 35 | 359.83 | 151.25 | 129.79 | 699.12 |
| BHB (mmol/L) | 35 | 0.74 | 0.17 | 0.37 | 1.06 |
| Relative abundance |  |  |  |  |  |
| *Caviibacter* | 35 | 22.05 | 36.55 | 0.00 | 99.81 |
| *Fusobacterium* | 35 | 6.54 | 11.08 | 0.00 | 49.09 |
| *Histophilus* | 35 | 6.70 | 19.49 | 0.00 | 82.43 |
| *Mycoplasma* | 35 | 9.23 | 12.25 | 0.00 | 58.33 |
| *Ureaplasma* | 35 | 6.19 | 16.13 | 0.00 | 89.27 |
| *Porphyromonas* | 35 | 5.45 | 12.05 | 0.00 | 61.55 |
| *Bacteroides* | 35 | 3.43 | 5.07 | 0.00 | 20.97 |
| *Cutibacterium* | 35 | 5.20 | 7.18 | 0.00 | 26.39 |
| *Staphylococcus* | 35 | 3.29 | 6.62 | 0.00 | 35.93 |
| *Streptococcus* | 35 | 3.33 | 13.66 | 0.00 | 81.38 |

<sup>1</sup>Average of all samples from disease diagnosis to 15 d postpartum.
<sup>2</sup>Average of all samples from 16 d postpartum until tissue collection at 1 mo postpartum.

**Table 3.** Results of the regression analysis for the effect of multiple regressor variables (Tables 2 and 5) on oocyte developmental competence for the Exp. 1 and Exp. 2.

| Study | No of obs. | Dependent variable | Regressor in the final model with P < 0.05 | Parameter estimate (slope) | Partial R-square | P-value |
| --- | --- | --- | --- | --- | --- | --- |
| Exp. 1 | 35 | Number of COC | *Cutibacterium* abundance | 0.49 ± 0.20 | 0.15 | 0.020 |
| Exp. 1 | 35 | Cleavage expressed as percentage of COC | *Fusobacterium* abundance | -0.64 ± 0.32 | 0.11 | 0.049 |
| Exp. 1 | 35 | Development to blastocyst expressed as percentage of COC | None at P < 0.05 | n/a | n/a | n/a |
| Exp. 1 | 35 | Development to blastocyst expressed as a percentage of cleaved embryos | *Mycoplasma* abundance | -0.58 ± 0.22 | 0.13 | 0.034 |
|  |  |  | *Cutibacterium* abundance | 0.85 ± 0.37 | 0.12 | 0.028 |
| Exp. 2 | 19 | Number of COC | None at P < 0.05 | n/a | n/a | n/a |
| Exp. 2 | 19 | Cleavage expressed as percentage of COC | *Ureaplasma* | -0.97 ± 0.28 | 0.28 | 0.004 |
|  |  |  | Late BHB | 144.1 ± 58.6 | 0.20 | 0.026 |
| Exp. 2 | 19 | Development to blastocyst expressed as percentage of COC | None at P < 0.05 | n/a | n/a | n/a |
| Exp. 2 | 19 | Development to blastocyst expressed as a percentage of cleaved embryos | None at P < 0.05 | n/a | n/a | n/a |

### 3.4 Predictive modeling of individual cow factors affecting in vitro embryo development for oocytes collected at 80 or 165 d postpartum (Exp. 2)

Plasma concentrations of haptoglobin, glucose, NEFA, and BHB are presented for cows based on their initial disease diagnosis and based on their disease status at 4 to 5 wk postpartum (**Table 4**). As expected, there was an effect (*P =* 0.044) of the initial disease diagnosis on plasma haptoglobin concentrations during the early period (greater in metritis cows; **Table 4**). Glucose (*P =* 0.282), NEFA (*P =* 0.376), and BHB (*P =* 0.431) were similar for healthy versus metritis during the early period. During the late period, plasma haptoglobin concentrations had decreased in metritis cows and were not different from healthy (*P =* 0.184; **Table 4**). Plasma glucose (*P =* 0.130), NEFA (*P =* 0.201), and BHB (*P =* 0.891) concentrations were similar for healthy compared with metritis cows.

**Table 4.** Plasma haptoglobin, glucose, non-esterified fatty acid (NEFA), and β-hydroxybutyrate (BHB) concentrations for Exp. 2. Cows were diagnosed as healthy or metritis during wk 1 to 2 postpartum. The vaginal microbiome at 4 to 5 wk postpartum was classified as either non-diseased or diseased as described [12]. Data are presented as mean ± SEM.

|  | Initial disease diagnosis |  |  | Disease status based on vaginal microbiome<br>at 4 to 5 wk postpartum<sup>1</sup> |  |  |
| --- | --- | --- | --- | --- | --- | --- |
|  | Healthy<br>(n=9) | Metritis<br>(n=10) | P-value | Non-diseased<br>(n=9) | Diseased<br>(n=10) | P-value |
| Early<sup>2</sup> |  |  |  |  |  |  |
| Haptoglobin (g/L) | 0.2 ± 0.2 | 0.8 ± 0.2 | 0.044 | 0.4 ± 0.2 | 0.6 ± 0.2 | ns |
| Glucose (mg/dL) | 50.4 ± 3.7 | 56.0 ± 3.5 | ns<sup>4</sup> | 51.9 ± 3.8 | 54.7 ± 3.6 | ns |
| NEFA (μEq/L) | 390.0 ± 62.9 | 311.3 ± 59.7 | ns | 402.7 ± 61.8 | 299.9 ± 58.6 | ns |
| BHB (mmol/L) | 0.84 ± 0.09 | 0.74 ± 0.08 | ns | 0.81 ± 0.09 | 0.77 ± 0.09 | ns |
| Late<sup>3</sup> |  |  |  |  |  |  |
| Haptoglobin (g/L) | 0.1 ± 0.1 | 0.2 ± 0.1 | ns | 0.1 ± 0.1 | 0.2 ± 0.1 | ns |
| Glucose (mg/dL) | 49.4 ± 3.9 | 58.0 ± 3.7 | ns | 46.5 ± 3.4 | 60.5 ± 3.2 | 0.008 |
| NEFA (μEq/L) | 247.1 ± 30.3 | 191.6 ± 28.7 | ns | 249.0 ± 30.1 | 189.9 ± 28.5 | ns |
| BHB (mmol/L) | 0.80 ± 0.04 | 0.82 ± 0.03 | ns | 0.81 ± 0.04 | 0.81 ± 0.03 | ns |

<sup>1</sup>Cows classified based on vaginal microbiome at 4 to 5 wk postpartum as described [12].
<sup>2</sup>Average of all samples from disease diagnosis to 15 d postpartum.
<sup>3</sup>Average of all samples from 16 d postpartum until 5 wk postpartum.
<sup>4</sup>ns = not significant.

None of the plasma metabolite concentrations during the early period differed for cows defined as non-diseased or diseased based on the vaginal microbiome at 4 to 5 wk postpartum (**Table 4**). Plasma concentrations of haptoglobin (*P =* 0.246), NEFA (*P =* 0.173), and BHB (*P =* 0.812) were similar for non-diseased versus diseased cows during the late sampling period (**Table 4**). Plasma glucose concentrations during the later period were less in cows defined as non-diseased (versus diseased) (*P <* 0.008; **Table 4**).

The average plasma concentrations of haptoglobin, glucose, NEFA, and BHB during the early and late periods as well as the relative abundance of the 10 most-abundant genera sequenced from the vagina at 4 to 5 wk postpartum of individual cows in Exp. 2 (**Table 5**) were included in a stepwise regression analysis (PROC REG of SAS) to build predictive models using a similar approach to that described for Exp. 1. Beginning with the 18 individual regressors (**Table 5**), the stepwise regression retained two regressors for the cleavage percentage [*Ureaplasma* abundance (negatively associated) and BHB during the later period (positively associated)] (**Table 5**). None of the regressors were significant for the number of COC, percentage of blastocysts developing from putative zygotes and the percentage of blastocysts developing from cleaved embryos (**Table 5**).

**Table 5.** Statistical parameters for individual regressor variables included in the PROC REG model for the prediction of oocyte developmental competence (Exp. 2).

|  | N | Mean | SD | Min | Max |
| --- | --- | --- | --- | --- | --- |
| Early<sup>1</sup> |  |  |  |  |  |
| Haptoglobin (g/L) | 19 | 0.50 | 0.69 | 0.01 | 2.62 |
| Glucose (mg/dL) | 19 | 53.34 | 11.06 | 38.16 | 88.65 |
| NEFA (μEq/L) | 19 | 348.57 | 187.71 | 29.60 | 645.83 |
| BHB (mmol/L) | 19 | 0.79 | 0.27 | 0.33 | 1.26 |
| Late<sup>2</sup> |  |  |  |  |  |
| Haptoglobin (g/L) | 19 | 0.13 | 0.34 | 0.01 | 1.52 |
| Glucose (mg/dL) | 19 | 53.90 | 12.20 | 37.12 | 90.09 |
| NEFA (μEq/L) | 19 | 217.91 | 92.80 | 79.17 | 458.06 |
| BHB (mmol/L) | 19 | 0.81 | 0.10 | 0.59 | 0.98 |
| Relative abundance |  |  |  |  |  |
| *Ureaplasma* | 19 | 13.48 | 21.69 | 0.01 | 78.76 |
| *Histophilus* | 19 | 9.53 | 14.75 | 0.00 | 47.51 |
| *UCG_005* | 19 | 7.42 | 5.70 | 0.05 | 16.33 |
| *Fusobacterium* | 19 | 5.63 | 11.37 | 0.00 | 44.33 |
| *Streptobacillus* | 19 | 5.60 | 9.54 | 0.00 | 34.73 |
| *Bacteroides* | 19 | 3.81 | 3.22 | 0.14 | 14.51 |
| *UCG_010* | 19 | 3.12 | 3.17 | 0.00 | 9.90 |
| *Mycoplasma* | 19 | 2.07 | 2.97 | 0.00 | 9.92 |
| *Rikenellaceae_RC9_gut* | 19 | 1.73 | 1.47 | 0.01 | 4.41 |
| *Caviibacter* | 19 | 1.69 | 5.03 | 0.00 | 19.63 |

<sup>1</sup>Average of all samples from disease diagnosis to 15 d postpartum.
<sup>2</sup>Average of all samples from 16 d postpartum until 5 wk postpartum.

## 4 Discussion

We hypothesized that uterine disease early postpartum would negatively affect oocyte developmental competence. Although this concept is well-established in the bovine literature, none of the supportive work included the collection and in vitro culture of oocytes from postpartum cows with a clinical diagnosis of healthy or with metritis. Results from the two experiments supported the premise that uterine disease can affect the developmental capacity of the oocyte and were consistent with respect to two important and novel findings. First, the original disease diagnosis that we made at 7 to 10 d postpartum did not affect the number of COC collected or their subsequent development when the oocytes were collected at approximately 1 mo postpartum (Exp. 1) or later postpartum (80 to 165 d, Exp. 2). The status of the uterus at approximately 1 mo postpartum, however, affected the number of COC that cleaved in vitro. Specifically, cows that had purulent material in the uterine lumen at 1 mo postpartum (Exp. 1) or cows that had a disease-associated vaginal microbiome at 4 to 5 wk postpartum (Exp. 2; d 80) had a lesser cleavage rate compared with non-diseased cows.

The observation that the disease status of the uterus at disease diagnosis (7 to 10 d postpartum) is less-important than the ultimate disease status of the uterus later postpartum is similar to what we observed in other studies of these two groups of cows [12,17]. We find that approximately one-third of cows that are diagnosed with metritis will recover within one month and will not develop endometritis (defined herein as purulent material in the uterine lumen).

Conversely, approximately one-third of cows that are diagnosed as healthy at 7 to 10 d postpartum will ultimately develop endometritis at one month postpartum. In our studies, the uterine architecture as well as uterine gene expression was more dependent on the diagnosis at 1 mo postpartum than the earlier disease diagnosis (7 to 10 d postpartum; [12,17]). Our observations are consistent with the results of Figueiredo et al. (2021) who concluded that failure of clinical cure following metritis ultimately determined fertility more so than the original diagnosis or treatment [18]

These observations suggest that the oocyte is sensitive to the extended period of uterine disease that defines endometritis. Systemic factors including bacterial endotoxins associated with endometritis are either sensed by the follicular cells or the oocyte itself and damage the developmental competence of the oocyte. The best-characterized of these are the lipopolysaccharides (LPS) that have been shown to change gene expression in follicular cells and inhibit the developmental competence of oocytes in vitro [19–23]. The inhibition of oocyte function does not necessarily arise from the uterus as poor udder health postpartum is associated with a reduction in fertility [24–26]. Our results may be applicable to the general finding that postpartum infectious diseases of cattle lead to subfertility [27], in part through an effect on the developmental competence of the oocyte.

All cows in Exp. 1 were sacrificed at one-month postpartum (single time point). Experiment 2 was smaller (fewer cows) and there was an early (approximately d 80) and later (approximately d 165) slaughter date. Nonetheless, we found the same effect (reduced cleavage rate) for oocytes collected but this effect was manifest at d 80 but not d 165. These data should be considered preliminary given the overall size of the experiment, but they are consistent with what is known about oocyte development. Specifically, oocytes require several months to develop from the primordial follicle stage [28] and a variety of insults during the developmental period (negative energy balance, heat stress, disease, etc. [6,29,30]) can compromise their developmental capacity and fertility. Our interpretation is that oocytes collected from cows at approximately 80 d postpartum (Exp. 2) had initiated their development at approximately 1 mo postpartum and were affected by the presence of uterine disease at that time. Oocytes collected at 165 d postpartum (Exp. 2) had not initiated their development while there was disease present in the uterus.

The results of the present study extend and generally confirm the relationship between uterine disease and oocyte development as reported in two other studies. In the first study, Ribeiro et al. [10] collected embryos from postpartum dairy cows with uterine tract disease versus no uterine tract disease. They showed an effect of uterine tract disease on the total number of cleaved embryos as well as live and high-quality embryos. In the second study, Dickson et al. [9] used nonlactating cows and an induced uterine infection model (infusion of pathogenic bacteria) to study oocyte development following transvaginal aspiration. They showed increased cleavage rate 2 d after the infusion but no effect on cleavage rate thereafter. They did show an effect on development to the morula stage that was manifest 24 d after the infusion. We failed to show any effect beyond the initial cleavage of oocytes in either Exp. 1 or 2. Neither study was identical to what was done here in that we used postpartum cows, all of whom were primiparous with a metritis diagnosis at 7 to 10 d postpartum with a subsequent diagnosis of endometritis (Exp. 1) or disease-associated vaginal microbiome (Exp. 2). Regardless, the general conclusion that uterine disease can affect embryonic development from fertilization onward held.

In addition to evaluating overt clinical signs of disease, we assessed the influence of systemic factors by measuring concentrations of an acute-phase protein (haptoglobin) and energy metabolites (glucose, NEFA, and BHB) during the postpartum period. These measurements were then analyzed in relation to the subsequent developmental competence of the oocytes. The association of bacterial genera within the reproductive tract with oocyte development in vitro was also evaluated. For these analyses, we used simple regression and a stepwise regression procedure to build a statistical model. These tests were performed to address the possibility that large shifts in circulating metabolite concentrations at the start of lactation can affect oocyte developmental competence. This concept is well-supported in the literature [6,30]. We used average concentrations of haptoglobin (an acute phase protein indicative of systemic inflammation [31]) and metabolites either before or after d 15 postpartum to bracket the periods of the most extreme changes in energy metabolites [32]. The approach was successful in capturing a range of metabolite concentrations for individual cows (**Tables 2** and **5**). The relative abundance of the most-abundance genera in the uterus (Exp. 1) or vagina (Exp. 2) as regressors was included as well to determine if a single or multiple species could explain the variation in oocyte development that we observed. The criteria for entry into the model was liberal (*P =* 0.15) but a regressor had to be significant (*P <* 0.05) to stay in the final model. Two important conclusions can be drawn from this analysis. First, despite including many regressors (**Tables 2** and **5**), we found very few associations with the number of oocytes or their subsequent development. The associations that were found were generally weak (r^2^ of 10 to 20%; **Table 3**), thus explaining a relatively small percentage of the total variation in the dependent variables that we measured (total number of COC, cleavage rate, etc.). There was only one example of a plasma metabolite concentration associated with embryo development (late postpartum BHB concentrations on cleavage rate). Most of the associations were with specific genera found in the uterine microbiome.

For Exp. 1, the relative abundance of the genus *Cutibacterium* was positively associated with both the number of COC and the development of cleaved embryos to blastocysts (**Table 3**). The genera is described as a commensal organism of the skin (specifically *Cutibacterium acnes*) but nonetheless we typically observe this genera in our 16S rRNA gene sequencing as well as bacterial culture studies [11,12,33,34]. The abundance of *Cutibacterium* averaged 5.2% within the uterus at 1 month postpartum (**Table 2**). *Cutibacterium* is typically not associated with bovine metritis or endometritis and was one of the genera that we observed increased at 7 to 10 d postpartum in healthy bovine uterus [11] and was cultured from uteri of slaughtered cows in Exp. 1 [11] and Exp. 2 [12]. In contrast to the positive associations with *Cutibacterium*, we noted negative association of embryo development with *Fusobacterium* and *Mycoplasma*.

*Fusobacterium* is one of the dominant uterine pathogens associated with bovine uterine disease [35] and we and others find abundant *Fusobacterium* in cows with metritis or endometritis [2,11,12,36]. The negative association of *Fusobacterium* abundance with embryo cleavage (**Table 3**) suggests that *Fusobacterium* infection can affect oocyte competence perhaps through a mechanism involving bacterial products entering the blood stream. Jeon et al. [36], for example, treated metritis cows with ceftiofur and reduced the abundance of *Fusobacterium* and LPS biosynthesis. The negative relationship between *Fusobacterium* abundance and embryo cleavage, therefore, is possibly explained by LPS from *Fusobacterium* inhibiting oocyte developmental competence. The second bacterial genera negatively affecting development was *Mycoplasma* (negative association with the percentage of blastocysts developing from cleaved embryos; **Table 3**). The primary mycoplasma species identified by sequencing were *Candidatus Mycoplasma*, *Mycoplasma wenyonii,* and *Mycoplasma bovigenitalium,* but none of these species explained a large percentage of the variation in the percentage of blastocysts from cleaved embryos. Nonetheless, each has been associated with infertility in the bovine either through infection of the reproductive tract (*Mycoplasma bovigenitalium* [37,38]) or systemic infection (hemotropic mycoplasmas; *Candidatus Mycoplasma* and *Mycoplasma wenyonii*) [39].

There were no significant regressors for number of COC or development of blastocysts in Exp. 2. The percentage of embryos that cleaved, however, was associated with the relative abundance of the genus *Ureaplasma* (negative association) as well as the circulating concentrations of BHB (positive association) from 16 to 30 d postpartum (late BHB). All the *Ureaplasma* sequenced in Exp. 2 was *Ureaplasma diversum,* an important species associated with infection of the reproductive tract of cattle [40,41]. *Ureaplasma* was the most-abundant genera that was sequenced from the reproductive tract of cows in Exp. 2 (**Table 5**). Both *Ureaplasma* and *Mycoplasma* belong to the family Mycoplasmataceae and lack a cell wall. Although the association with in vitro embryonic development differed (*Mycoplasma* affecting blastocyst development in Exp. 1 and *Ureaplasma* affecting embryo cleavage in Exp. 2), the observation that closely related bacterial species are impacting in vitro embryonic development across two different studies is worth noting and should be studied further. We did find in a previous study that there was greater *Ureaplasma* abundance in the uterine lumen of cows with endometritis (purulent material in the uterus [11]).

The early (first 15 d postpartum) haptoglobin concentrations were greater in metritis compared with healthy cows in Exp. 1 (**Table 1**) and Exp. 2 (**Table 4**). Other metabolite concentrations were inconsistent across studies with respect to disease status either at disease diagnosis (7 to 10 d postpartum) or one-month/4 to 5 wk postpartum. The only metabolite concentration that was associated with embryo development in either Exp. 1 or 2 was the late BHB concentrations in Exp. 2 (**Table 3**; positive association with embryonic cleavage). In their study of metabolic parameters and oocyte quality, Matoba et al. [42] reported a positive association between oocyte cleavage and NEFA concentrations and this result is consistent with our observation with a positive association between BHB and embryonic cleavage. Nonetheless, the conclusion of the Matoba et al. [42] paper was that their data did not support an effect of lactation-induced metabolic stress on oocyte development. This conclusion is like the one that we make from Exp. 1 and 2. The failure to associate metabolite concentrations with oocyte developmental capacity is inconsistent with a large volume of scientific literature that suggests elevated NEFA and BHB are inhibitory to oocyte development when added to culture medium in vitro [6,30]. The present study was primarily addressing metritis and we did not attempt to manipulate metabolite concentrations through diet, etc. We also used first parity cows exclusively that are lower-producing and less-likely to have metabolic disease [43]. We did, however, have a considerable range in both early and late metabolite concentrations (**Tables 2** and **5**) and failed to identify associations with the number of oocytes that we collected or their development in vitro.

We recognize the limitations of this work. According to the 2022 summary of global statistics by the International Embryo Technology Society (https://www.iets.org), the percentage of transferable embryos from collected oocytes was 21.7% across all nations. In the current work, the percentage of blastocysts produced from individual oocytes was 14.0% (Exp. 1) and 17.3% (Exp. 2). The percentage of blastocysts across both experiments was approximately 5 percentage points below the industry average. A greater percentage of blastocysts across both experiments may have improved the sensitivity of our analyses for blastocyst development.

Cows in Exp. 1 were sacrificed and their uterine horns flushed to classify each cow as either purulent or clear. These definitions were then used to classify the status of individual cows at 1 mo postpartum. Cows in Exp. 2 were not sacrificed at 1 mo postpartum nor was the uterus sampled to measure the microbiome in the uterine lumen or whether the cows had endometritis. The primary objective of Exp. 2 was to study the uterine microbiome at later stages postpartum following slaughter. As such, we did not want to introduce swabs or a cytobrush (cytological evaluation [44]) into the uterine lumen at any time before slaughter (d 80 to d 165 postpartum).

We used the vaginal microbiome during the first 5 wk postpartum to classify the microbiome of individual cows using PC analyses into either non-diseased or diseased states. This classification assumed that the vaginal microbiome reflects the uterine microbiome. In a previous study [12], we demonstrated that this approach could identify significant differences in differential gene expression in the uterus of non-diseased and diseased cows at the time of slaughter [12]. Here, we applied the same classification and observed that it also correlated with differences in the percentage of cleaved embryos. There are clear advantages of a specific diagnosis of endometritis at one month postpartum, and we concede that the approach we used had limitations.

In conclusion, we examined the capacity of oocytes to develop in vitro when collected from healthy or diseased dairy cows at approximately 1 mo (Exp. 1) or 80 to 165 d (Exp. 2) postpartum. Across both studies, we noted that the percentage of cleaved oocytes was decreased for cows classified as diseased based on purulent contents of the uterine lumen (Exp. 1) or vaginal microbiome (Exp. 2). We also found in Exp. 2 that the sensitivity of the oocyte to uterine disease was present at 80 d but not 165 d postpartum. We did not find a strong association between circulating metabolite concentrations and measurements of embryo development in vitro. The development of oocytes was positively associated with *Cutibacterium* abundance, and negatively associated with the abundance of *Fusobacterium*, *Mycoplasma*, and *Ureaplasma* in the reproductive tract. These studies extend the findings of related work showing the impact of disease on the capacity of oocytes to develop following fertilization. The reduced developmental capacity of oocytes from diseased cows may partially explain the decreased fertility observed in cows with uterine or other inflammatory diseases postpartum.

## Credit authorship contribution statement

**Monica Caldeira:** Investigation, Formal analysis, Writing - original draft preparation, reviewing and editing. **Katy McDonald:** Investigation. **Ethel Martinez:** Investigation. **Joao Moraes:** Investigation, Writing – Review & Editing. **Isabella Sellmer Ramos:** Investigation. **Scott Poock:** Investigation. **M. Sofia Ortega:** Supervision, Methodology, Resources, Investigation, Writing – Review & Editing. **Matthew Lucy:** Conceptualization, Funding acquisition, Supervision, Writing – Review & Editing.

## Acknowledgments

The authors would like to express their gratitude to the employees of Foremost Dairy Research Center (Columbia, MO), and Ohlde Dairy (Lynn, KS) for their assistance in the management of cows.

